# Microfluidic device combining hydrodynamic and dielectrophoretic trapping for the controlled contact between single micro-sized objects and application to adhesion assays

**DOI:** 10.1101/2022.11.08.515593

**Authors:** Clémentine Lipp, Laure Koebel, Romain Loyon, Aude Bolopion, Laurie Spehner, Michaël Gauthier, Christophe Borg, Arnaud Bertsch, Philippe Renaud

## Abstract

The understanding of cell-cell and cell-matrix interactions via receptor and ligand binding relies on our ability to study the very first events of their contact. Of particular interest is the interaction between a T cell receptor and its cognate peptide-major histocompatibility complex. Indeed analyzing their binding kinetics and cellular avidity in large-scale low-cost and fast cell sorting would largely facilitate the access to cell-based cancer immunotherapies. We thus propose a microfluidic tool able to independently control two types of micro-sized objects, put them in contact for a defined time and probe their adhesion state. The device consists in hydrodynamic traps holding the first type of cells from below against the fluid flow, and a dielectrophoretic system to force the second type of object to remain in contact to the first one. First the device is validated by performing an adhesion frequency assay between fibroblasts and fibronectin coated bead. Then, a study is conducted on the modification of the cellular environment to match the dielectrophoretic technology requirements without modifying the cells viability and interaction functionalities. Finally, we demonstrate the capability of the developed device to put cancer cells and a population of T cells in contact and show the discrimination between specific and non specific interactions based on pairs lifetime.

## 1. Introduction

Multicellular organisms require a very well organized and finely balanced cell-cell communication, adhesion, coordination and programmed cell death to ensure the organism’s homeostasis. These functions rely on specialized receptors placed at the cells membrane whose binding to their ligand triggers a defined function. Receptor-ligand interaction is thus a cornerstone of multicellularity, not only maintaining the cells physically adhered to one another, but also enabling communication to ensure function and organism homeostasis. Malfunction of the receptors to properly trigger the appropriate response leads to imbalances in the organism and failure to exert specific functions and our understanding of these pathologies thus relies on the capacity to study such interactions. Because they characterize the very first instant of the reaction chain induced by a binding event, the binding kinetics parameters defining the rate of bond formation and dissociation are of particular interest. Receptors whose ligand are in solution benefit from well characterized tools to study their binding kinetics, such as surface plasmon resonance (1). In contrast, tools available to study receptors whose ligand are anchored on a cell or on a surface remain relatively inaccessible.

Measuring the binding kinetics of two surface anchored receptors and ligands demands the independent control on the spatial position of two micrometer sized objects with high temporal resolution, which is technically challenging (2). State of the art methods for such assays comprise atomic force microscopy (AFM), which is a precision tool of choice to measure the force characteristics. Typically, a cell is picked and attached to the tip of a soft microfabricated cantilever using either surface functionalization (3) or an embedded fluidic system (4) and used to contact another cell adhered to a substrate. The time of contact can be controlled before probing the adhesion state and measuring the force resistance of the formed bond(s) through the deformation of the cantilever. Single bond detachment can be resolved thanks to the high force sensitivity of the probe. Alternatively, the dual pipette assay (DPA) makes use of two micropipettes to attach one cell per pipette via the aspiration of the cells. The pipettes can then be guided using micromanipulators coupled to high resolution high speed imaging to probe the adhesion state as a function of time of contact (5). Variations of this method comprise the attachment of a red blood cell coated with a specific ligand to one of the cells. Indeed red blood cells are soft and a precise force measurement of the bond can be deduced from their deformation. Alternatively, a bead coated with the ligand can be attached to an aspirated red blood cell and put in contact to the receptor presenting cell attached to the other pipette. In that case, the thermal fluctuations of the bead are measurable because of the high deformability of the red blood cells and indicate the adhesion state of the pair (6). A direct measurement of the on and off rates from the adhesion state timeline is then made possible. Optical tweezers (OT) are another tool that makes possible the precise manipulation of objects thanks to the attraction of particles towards the waist of a highly focused laser beam. OT are thus able to both push an object trapped in the waist towards a cell adhered to a surface or trapped in another OT as well as pull on it with known force to probe the adhesion state and force of adhesion (7,8). These three methods rely on a very soft spring linked to one of the cells to push and pull it to and from another cell to allow a controlled force and time of interaction. They are thus highly precise in force and position control but are also of very low throughput and require highly skilled staff to perform the assay.

Thanks to their ability to control position of micro-sized objects through various forces, microtechnologies have a potential to control the contact between two objects at larger throughput than the above mentioned methods. The strategy of confining two objects in the same micro-sized environment was largely used to study downstream effects after the contact (9–11), but lacks the possibility of independently controlling the two objects to study binding kinetics. Systems for the dynamic and serial cell-cell and cell-bead interaction were developed in microfluidic chips by immobilizing in a first step the first type of particle using monolayer adhesion (12,13), hydrodynamic traps (14,15) and hydrodynamic traps combined with sedimentation (16). The second type of cell was flown on top and reduction in speed upon contact between the two objects, pair lifetime, resistance to flow shear stress or downstream signalling molecules monitored to characterize the interaction. These systems are powerful tools to probe receptor-ligand interactions at higher throughput than the macrosized methods presented above, but they lack a possibility to control both force and contact time between the two objects.

Controlled contact between cells is of specific interest for cell-based cancer immunotherapies. Indeed the process of selecting patient T cells with high antitumor activity is currently a long and very expensive process, limiting the generalization of such approach despite their proven efficacy (17). In this context, a tool able to reliably pair T lymphocytes with antigen presenting cells (APCs) or tumor cells and rapidly assess the specificity of the interaction could simplify the long process of T cell selection and facilitate in the long term the access to this kind of therapy. In this context, we propose a microfluidic tool to control cell-cell and cell-bead contact in a microfluidic chip using forces deriving from a different phenomenon for each object. The effects are thus orthogonal for an independent manipulation of the two micro-sized particles. The method relies on the combination of two previously published trapping methods based on planar hydrodynamic trapping of cells (18) and dielectrophoretic (DEP) trapping (19), both physical principles being widely used to precisely manipulate and trap single cells (20,21). We first present the working principle of the microfluidic device and its capabilities (Section). The functionality of the tool is then validated by performing an adhesion frequency assay with fibroblasts cells and fibronectin coated beads and the extracted binding kinetics parameters are compared to literature (Section). In order to demonstrate the potential of such a tool in immunotherapy applications, the pairing of T cells with cancer cells is then performed and the effect of the binding of T cells receptors (TCR) to its cognate peptide-major histocompatibility complex (pMHC) studied (Section). The measurement of the pair’s lifetime was measured and the effect of TCR-pMHC bonds in this latter are demonstrated. A measurement of cellular avidity based on these measurements is then proposed by assigning a pair lumped off rate as metric. The characteristic of the device are finally discussed in Section as well as future developments and perspectives.

## 2. Material and Methods

### 2.1. Microfabrication

Buried microchannels were fabricated as detailed by (18) for the planar hydrodynamic trapping. Shortly, 500 nm Al_2_O_3_ was sputtered on a fused silica substrate and access holes as well as traps and outlets were defined by photolithography and etched using ion beam etching. The channels are defined by exposure to a vapor phase of hydrofluoric acid (HF) that selectively underetches the fused silica substrate without attacking the Al_2_O_3_ layer. Access holes are then sealed by depositting a 2.5 μm thick layer of low temperature oxide doped with boron and phosphate (BPSG) leading to 3 μm diameter traps. Electrodes for the dielectrophoretic actuation are then defined by a liftoff process. The negative resist AZ nLOF (MicroChemicals) is coated (ACS 200, Süss), exposed (MLA 150, Heidelberg Instruments) and developed (ACS 200, Süss) on top of the structured fused silica substrate. 20 nm of titanium and 200 nm of platinum are evaporated (EVA 760, Alliance-Concept) and lifted off in a remover bath (MICROPOSIT Remover 1165). The remaining photoresist residues remaining in the buried channels are then cleaned using an oxygen plasma (GIGAbatch, PVA TePla). The top PDMS channels are then molded, aligned and permanently bonded to the electrodes and glass channels as described in (19). The full fabrication process is illustrated in Figure S4 in supplementary material, and pictures of the fabricated combined traps and the whole final chip respectively in Figures S5 and S2A.

### 2.2. Experimental platform and protocols

The chips were primed with Pierce protein-free (PBS) blocking buffer for 2 hours to prevent proteins from adhering to the surfaces. The cells or beads were placed in a chromatography vial connected to the punched PDMS with 360 μm outer diameter tubing for tight sealing. Pressure was applied to the vials using Fluigent Flow-EZ pressure controllers. The chip was mounted on and electrically connected to a custom PCB placed on the stage of a Leica DMI3000 B inverted microscope and observed using a uEye (IDS) camera. All the electric signals needed to control the positions of the particles are sent through a home made PCB creating the multiplication of an AC signal at 100 kHz and different DC signals whose amplitudes are controlled by the computer with an adapted C++ program through an analog output generator (Mccdaq USB-3100). The full setup is illustrated in Figure S1 and the chip and PCB mounting in Figure S2B in supplementary material.

The peak value for deviation voltages *V_1_* and *V_2_* was set to 10 V whereas the peak trapping voltages *V_sync_* and *V_contact_* were set to 8 V. Pressures during contact experiments were set to 15 mbar to ensure constant drag force exerted on the DMPs.

### 2.3. Cell culture and preparation

The adherent mouse fibroblast cell line NIH-3T3 was cultured in medium (DMEM, high glucose, GlutaMAX Supplement) supplemented with 10% FBS and 1% penicillin-streptomycin at 37 °C in a 5% CO_2_ atmosphere. The semi-adherent cancer cell line Colo205 was cultured in medium (RPMI) supplemented with 10% FBS and 1% penicillin–streptomycin at 37 °C in a 5% CO_2_ atmosphere. Colo205 tumor cell lines were pulsed with 10 μg/mL of specific or irrelevant peptide, and incubated for 1h at 37°C in 5% CO_2_ in RPMI medium supplemented with 10% FBS and 1% penicillin-streptomycin. The specificity of CD8+ T cells towards the peptide was previously assessed by co-culture of 5·10^4^ T cells with pulsed tumor cell lines in the presence of Golgi Plug (BD Biosciences) in 96-well plate at 1:1 E:T cell ratio.

Staining of the cells was performed by incubating the cells in PBS with 4 μm Calcein UltraBlue AM (Cayman Chemical) or with 1 μm Calcein AM (Invitrogen™) for 1 h. The working solution is composed of 40% RPMI and 60% deionized water. The solution is compensated for osmolarity by the addition of dextrose (Sigma-Aldrich) and cleaned through a 0.22 μm filter. After staining, the cells were resuspended in the working solution and passed through a 40 μm cell strainer before the experiment. All reagents are from Gibco unless specified.

### 2.4. Medium compatibility protocols

For cell viability assay, cells were immersed for 5 h in complete medium diluted in different percentage of DI water with corresponding amount of dextrose to compensate for osmolarity at 37 °C in a 5% CO_2_ atmosphere. After 5 h, the cells were resuspended in standard complete medium and cultivated at 37 °C in a 5% CO_2_ atmosphere for 48 h. Cell viability was assessed using trypan blue after the 5 h incubation in custom medium and 24 h and 48 h after resuspension in standard medium.

For cell activation assay, CD8+ T cells were stimulated with specific or irrelevant peptides at 5μg/mL (JPT Technologies) in the presence of Golgi Plug (BD Biosciences) in complete medium (RPMI medium supplemented with 10% FBS and 1% penicillin-streptomycin) or in complete medium diluted in different percentage of DI water with corresponding amount of dextrose to compensate for osmolarity. CD8+ T cells were incubated at 37°C in 5% CO2 for 5h for cell viability and cytokine production analysis. Cytokine production was analyzed by flow cytometry upon cell staining performed according to the manufacturer’s instructions (BD). Briefly, cells were stained for 30 min with Live/Dead fixable dead cell stain (eBioscience^TM^ Fixable Viability Dye eFluo^TM^ 506 Invitrogen), anti-CD3 (PB, BD Pharmingen), anti-CD4 (APC-H7, BD Pharmingen) and anti-CD8 (PE, Diaclone). For intracellular staining, cells were incubated with Cytofix/Cytoperm^TM^ (BD Biosciences) for 30 min then stained with anti IFNγ(APC, BD Pharmingen) and anti TNFα(FITC, BD Pharmingen). Cells were acquired on a BD FACS-Canto II (BD Biosciences) flow cytometer and data analyzed using FlowJo Software.

### 2.5. Bead coating

5 μm in diameter polystyrene beads coated with streptavidin were purchased from Spherotech. The beads were cleaned 3 times in PBS and resuspended in a solution of PBS with 20 μg/ml biotinylated fibronectin (Cytoskeleton) or biotinylated BSA (Pierce) and left to incubate for 1 h at room temperature and with gentle rotation. The beads were then washed three times in PBS supplemented with 7.5% BSA to prevent the beads from adhering to each other and resuspended in the working solution.

## 3. Results

### 3.1. Principle of operation: Combining hydrodynamic traps and dielectrophoresis in a microfluidic chip

We present a microfluidic system combining fluidic and dielectrophoretic actuation capable of performing in flow cell-particle interaction. The device is illustrated in Figure 1 a and comprises two inlets for medium perfusion controlled by the pressures *P_in_*,_1_ and *P_in_*,_2_ followed by serpentines channels to increase its flow resistance. The two channels merge before entering the interaction chamber. This latter is composed of an upstream DEP actuated deviation system capable of steering the incoming particles to specific streamlines (22), followed by an expansion of the channel and two lines of DEP traps. Each line is made of coplanar electrodes and comprises four distinct trapping units capable of focusing the particles to one point in three dimensions (19) as well as a bypass zone to which unwanted particles can be deviated to leave the chamber. The upstream line is named the synchronization line because its role is to synchronize the arrival of particles to the second line once the amount of particles per trap is reached. The second line is named the contact line and features hydrodynamic traps placed along the centers of each one of the DEP traps. The hydrodynamic traps are precisely described in (18) and feature holes, placed at the bottom of the main PDMS channel and connected to a second lower level of channels, named buried channels. The buried channels connect the traps to a pressure control channel, whose pressure *P_c_* defines the flow direction in the buried channel. Particles of larger diameter than the trap and lying in the streamlines passing through the traps and buried channels clog the traps due to their larger dimensions, stopping the flow in the buried channels and immobilizing the particle thanks to the pressure difference built across it. Figure 1 b is an optical microscope image of the synchronization (top) and contact (bottom) lines with transparent buried channels and hydrodynamic traps. Figure 1 d is a scanning electron microscopy image with two hydrodynamic traps surrounded by coplanar electrodes (highlighted in yellow for better visualization).

**Fig.1.**
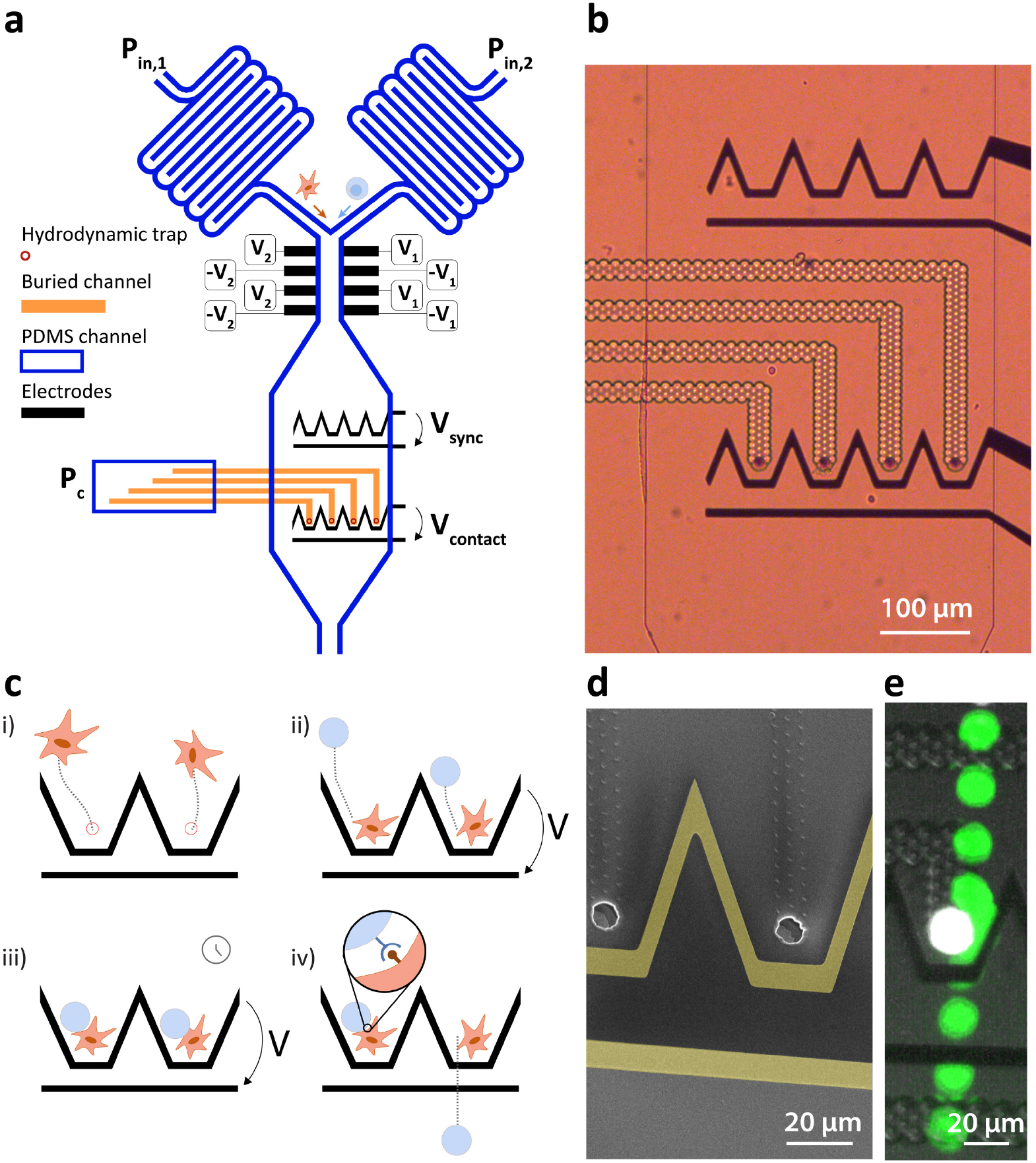
**a** Schematic representation of the microfluidic chip and electrodes with controlled voltages and pressures. The orange part represent the buried channel embedded in the glass substrate and running at a lower level under the PDMS channel. **b** Optical microscope picture of the interaction zone with the synchronization line (top) and contact line (bottom). Each funnel shaped electrode feature generates a DEP force field with single position of equilibrium against the flow at their center when a voltage is applied. The contact line comprises hydrodynamic traps at the center of the DEP traps. **c** Workflow steps for the controlled interaction between objects with emphasis on two traps of the synchronization line. **d** SEM picture of two units of the synchronization line. The hydrodynamic trap is visible at the center of the funnel shaped electrodes. **e** Timelapse image of the controlled contact between two cells. The hydrodynamically trapped cell is coloured in white and the dielectrophoretically manipulated cell in green by image processing for clarity.

The interaction zone thus comprises DEP and hydrodynamic traps, whose roles are to exert independent effects on the particles. Once a particle is immobilized in the hydrodynamic trap, the DEP force does not affect it anymore because it is weaker than the force exerted by the difference of pressure across the hydrodynamic trap. This particle is hereafter named the Hydrodynamically Trapped Particle (HTP). A second particle however is affected by the DEP induced force when a difference of potential is applied across the contact electrodes. Because the hydrodynamic trap is placed at the equilibrium position of the DEP trap, its effect directs the second particle, hereafter named Dielectrophoretically Manipulated Particle (DMP), towards the HTP and immobilizes it in close contact to the HTP against the flow. The workflow steps for the controlled interaction are depicted in Figure 1 c with emphasis on two traps of the contact line. The first step consists in the introduction of the first type of particles in the interaction chamber by increasing *P_in_*,_1_ and setting *P_in_*,_2_ to zero. The particles are then immobilized in the hydrodynamic traps by setting a negative value to *P_c_* (Figure 1 ci)). Once all the traps are filled with HTPs, *P_c_* can be increased to a value close to zero to minimize the difference of pressure across them and associated deformation. The second type of particles are then introduced in the chamber by increasing *P_in_*,_2_ and setting *P_in_*,_1_ to zero. The arriving particles are first steered towards each of the trapping units of the synchronization line (not shown) using the deviation system and trapped against the flow drag force under the effect of the DEP force generated by *V_sync_*. The destination of the particles is defined by the ratio of voltage applied to each side of the channel *V_1_/V_2_* (22). Once the synchronization line is filled by one particle per trap, *V_sync_* can be turned off and the DMPs are dragged downstream by the flow and immobilized in close contact to the HTP by turning on *V_contact_* (Figure 1 c ii)). The voltage is maintained for the desired contact duration (Figure 1 c iii)). Once this duration is elapsed, *V_contact_* is turned off and the DMPs are dragged away from the HTP by the constant flow. Adhesion events mediated through receptor-ligand bonds can be observed and related parameters such as state of adhesion or lifetime of the pairs can then be assessed and measured (Figure 1 c iv)).

Figure 1 e is a timelapse image of the contact between two cells demonstrating the device’s capability, the HTP is coloured in white and the DMP in green by image postprocessing for clarity.

### 3.2. Validation of the device via the interaction between fibronectin and fibroblasts

In order to validate the device, a well known and characterized pair of receptorligand interaction was studied using the developed microfluidic chip. Integrins present on the surface of mouse fibroblasts were shown to bind specifically to fibronectin coated on a surface (7). For the specific case of a species outnumbering the other one, the mathematical expression describing the probability of adhesion of one bond after a contact time *t* was derived (5):

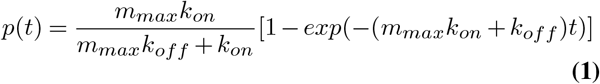

With *k_on_* and *k_off_* the forward and reverse binding constant in *μm^2^s*^-1^ and *s*^-1^ respectively and *m_max_* the surface density of the most abundant species out of the two in *μm*^-2^. The probability *p_n_* of forming *n* bonds after a time *t* is then given by the binomial distribution:

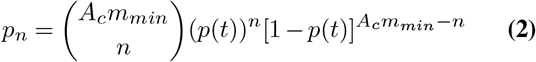

With *A_c_* the surface of contact in *μm*^2^ and *m_min_* the surface density of the less abundant species in *μm*^-2^ and 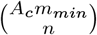 the binomial coefficient. The probability of adhesion *P_a_* mediated by minimum *n_min_* bonds is then given by:

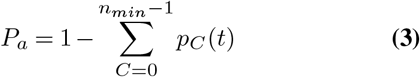

For cases where a single bond is sufficient to mediate an adhesion event, the last expression becomes:

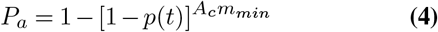

In this experiment, 5 μm in diameter polystyrene beads coated with fibronectin were taken as DMPs and were put in contact during a controlled amount of time to mouse fibroblasts who played the role of HTPs. The adhesion state was assessed after the forced contact was released. The negative control consisted of 5 μm in diameter polystyrene beads coated with bovine serum albumin (BSA) as DMPs. Figure 2 a illustrates a controlled contact of 3 seconds between a fibronectin coated bead and a fibroblast that does not display any adhesion after the contact. The measure was taken for more than 20 events per contact time and the fraction of adhesion events is reported as a function of contact duration in Figure 2. The contact time was measured from video analysis and taken as the time the bead was immobile in contact to the HTP. Disparities in contact duration were observed and its standard deviation is reported as the horizontal error bar. The large majority of beads had a position of equilibrium downstream of the HTP between this latter and the electrode, ensuring repeatable experimental conditions. The vertical error bar represents the 95% confidence interval on the percentage calculated from a binomial distribution of the adhesion events.

**Fig. 2.**
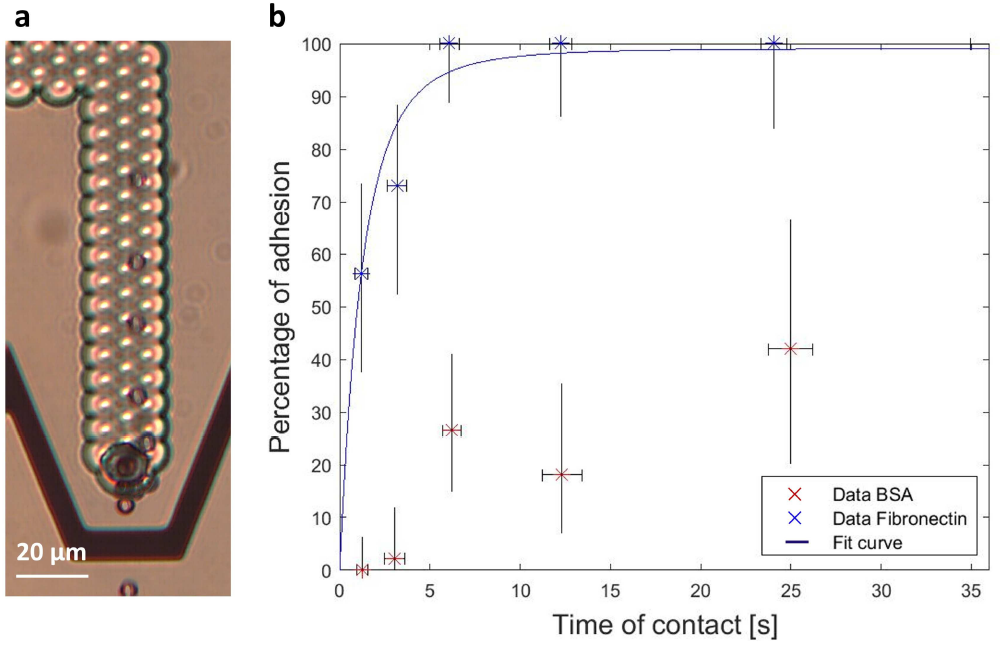
**a** Timelapse image of a fibronectin bead placed in contact to a mouse fibroblast during 3 seconds. No adhesion was observed after the contact and the bead is dragged downstream after the release. **b** Percentage of adhesion events as a function of contact duration for fibronectin coated beads as DMPs and mouse fibroblasts as HTPs. BSA coated beads as DMPs are used as a control. We demonstrate that the adhesion is mediated by integrin receptors at the surface of the fibroblasts binding specifically to fibronectin and extract the binding parameters from the fitting Equation 4 to the data.

Integrins were taken as the less abundant species and one bond considered as sufficient to mediate an adhesion as deduced by the similar work performed by Thoumine *et al*.(7). As expected from equation 4, the percentage of adhesion of fibronectin coated beads increases as a function of contact duration until reaching a plateau. The control data displays a significantly lower percentage of adhesion with no adhesion event reported for the lower range of contact duration. Experimentally, the strength of adhesion was significantly lower for BSA as the adhered beads would detach from the cell at very low flow rate, contrary to most fibronectin beads who remained attached even when increasing the flow rate until the maximum value allowed by the pump. Equation 4 was fitted to the data and is represented as the continuous line on the graph of Figure2 b. The parameters extracted from the best fit are *k_r_* = 0.17 *s*^−1^ and *A_c_m_min_m_max_k_f_* = 0.76 *s*^-1^. This latter parameter is constant for a given adhesion curve and was found to be in the range reported by the similar adhesion experiment by Thoumine *et al*.(7).

### 3.3. Application to the measurement of T cell - cancer cell pairs lifetime

The device was used to test more complex receptor-ligand pairs such as TCR-pMHC. An HLA-A2-restricted clone of human CD8+ T cells was thus used as DMPs and cancer cells pulsed with the peptide to which the TCRs of the T cells clone are specific were used as HTPs. The negative control consisted of the same cancer cells pulsed with an irrelevant peptide.

#### 3.3.1. Medium compatibility

Because the presence of an electric field in a conductive medium induces movement of ions and ensuing Joule heating effect (23), diluted media with lower conductivity must be used to manipulate particles with DEP. Typical medium for DEP manipulation is composed of 10% PBS and 90% deionized (DI) water supplemented with dextrose to correct for osmolarity (24). T cells however need specific medium compounds to properly activate following a specific binding of their TCR (25). Furthermore, prolonged exposition to modified media can compromise their viability, which is necessary for further culture after selection in cell based immunotherapies. The maximum time necessary for T cell selection in a chip was estimated to 5 h and the activation and viability of the T cell clones in different dilutions of medium in DI water and corrected for osmolarity by the addition of dextrose after incubation with specific peptide was thus assessed using the protocol described in the Experimental Section. The T cell clones immersed in 10% PBS not only did not produce IFN-γ and TNF-α after co-incubation with the specific peptide but only 8% of the initial cells remained alive after 5 h immersion in the specific medium and 48h re-culture in standard medium (Supplementary Figure S6). Activation and viability tests after a 5 h exposition to the medium were thus performed in different dilutions of RPMI supplemented with FBS and the results are presented in Figure 3 and Supplementary Figure S7 indicating a minimum standard medium content of 40% to ensure proper activation of the clones as well as viability after immersion during 5 h. This concentration was thus selected to perform the on-chip interaction experiments.

**Fig. 3.**
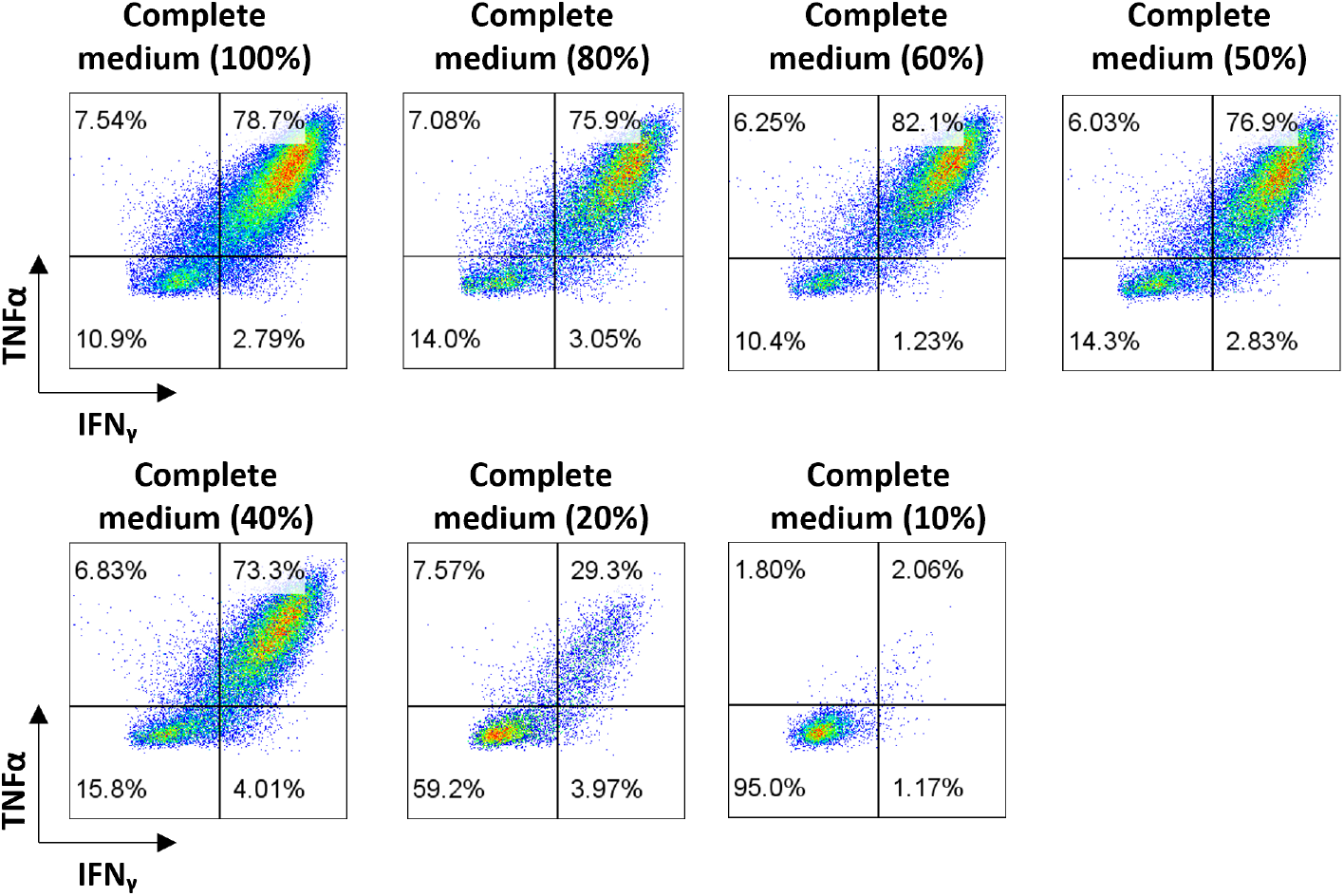
Intracellular IFN-**γ** and TNF-**α** content after 5 h co-incubation of T cell clones with specific peptide in medium composed of different percentage of RPMI supplemented with FBS and diluted in DI water with the adapted amount of dextrose to compensate for osmolarity. A minimum content of 40% complete medium was found to be necessary to ensure T cell clones activation and viability.

#### 3.3.2. Pairs lifetime results

T cells and cancer cells were stained with calcein AM of distinct color (blue and green respectively) to be able to differentiate between cell types, but also to visualize any event of electroporation or membrane damage due to the difference of pressure applied by the hydrodynamic traps. Indeed, calcein is a small molecule without covalent binding to intracellular compounds that quickly diffuses out of the cell in case of membrane damage. Only events with intact cells displaying constant and bright fluorescence were considered for the analysis. The average duration of contact was measured as the time the cells were immobile and in contact to the HTPs and is of 22 seconds with standard deviation of 3 seconds. The large majority of T cells had a position of equilibrium downstream of the HTPs between this latter and the electrode, ensuring repeatable experimental conditions. The pair lifetime was defined as the time a T cell stays in a radius of 20% that of the cancer cell once the voltage of the interaction line *V_contact_* is turned off. A picture of a forced contact between the two cells is illustrated in Figure S3 and a video of a contact provided in Supplementary video SV1. The results of the lifetime measurements are displayed in Figure 4 a. Adhesions were observed in both control and tested case, indicating that the expected TCR-pMHC bonds are not the only receptor-ligand pairs responsible for adhesion. Indeed T cells have different receptors responsible for rolling on the endothelium, cell migration towards their

**Fig. 4.**
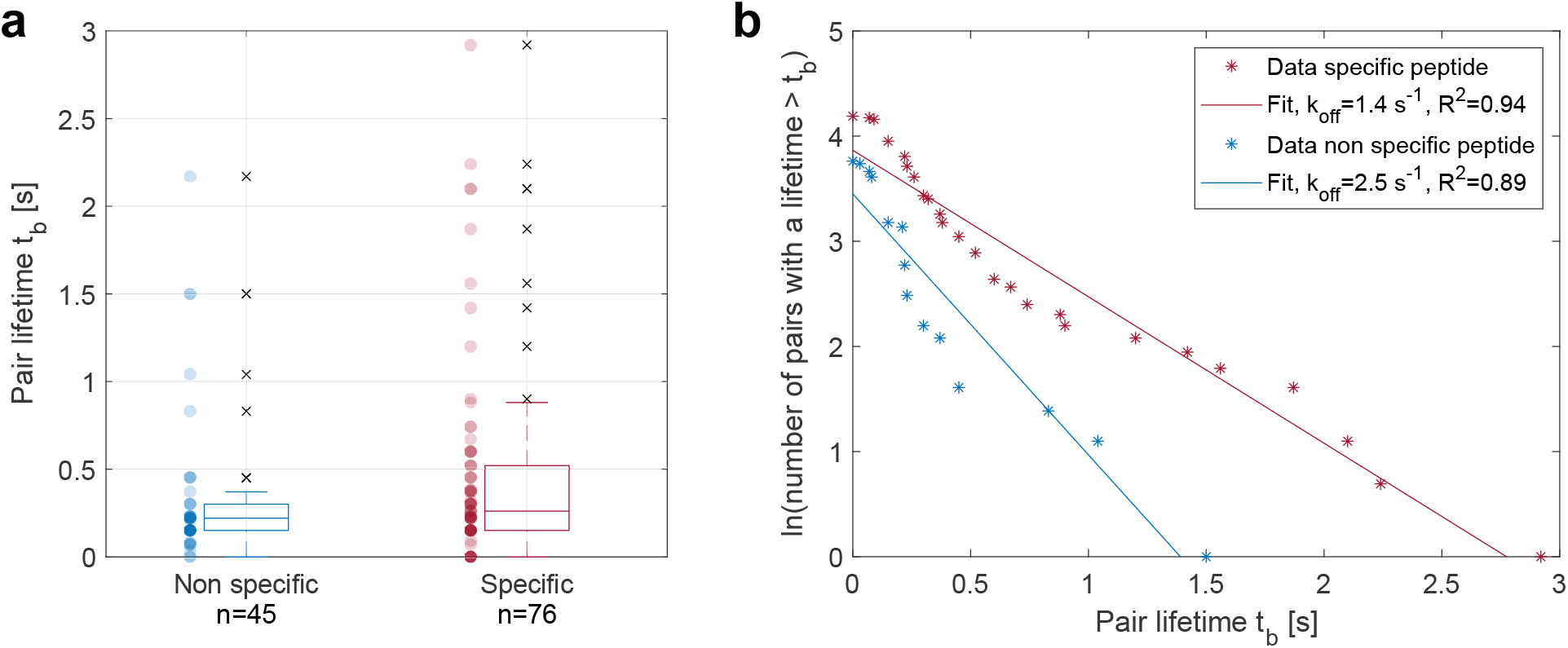
**a** Pair lifetime measurements after 22 seconds of forced contact between Colo205 and peptide-specific T cell clones in the case where the HLA-A2+ Colo205 cells were pulsed with the TCRs cognate peptide (specific) or irrelevant peptide (non specific). **b** A lumped off rate can be fitted to the data indicating a faster dissociation of pairs without specific peptide than the pairs with specific peptide.

targets and mediating the immune synapse such as LFA1-ICAM, CD28-CD80/CD86 or CD2-CD48/CD59 (26).

The samples distributions were tested for normality using a Kolmogorov-Smirnov test and the null hypothesis of a normal distribution was rejected is both cases. The samples were thus tested for equal median using Mann-Whitney U test and obtained a p-value of 0.03, rejecting at 95% confidence interval the null hypothesis and indicating a larger median lifetime in the presence of specific peptide on the cancer cells. The TCR-pMHC interaction thus tends to increase the pairs lifetime.

Lifetime of single bonds following first order dissociation kinetics can be described by an exponential decay and the probability *P_i_* of a bond formed at time *t* = 0 to remain intact at time *t_b_* follows the law:

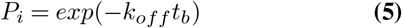

As discussed above, the adhesion events observed are not only mediated by multiple bonds but also by different receptor-ligand pair types. Avidity represents best the present situation than affinity as it is defined as the strength of an interaction between cells mediated by multiple receptor-ligands such as the TCR, the co-receptor CD8, other adhesion molecules and activating/inhibitory molecules. It is mainly measured via multimer binding assays and is believed to be a better predictor of T cell effector function than simple TCR-pMHC affinity (27,28). We thus approximated the cell-cell dissociation rate to a first order dissociation kinetics and fitted the results to extract a lumped off rate characterizing the avidity. Figure 4 b presents the natural logarithm of the number of events with a lifetime longer than *t_b_* as a function of time and is thus a representation of *In(P_i_(t))*. A linear dependency was fitted to the data and the slope taken as the effective off rate, which was measured to be of 1.4 *s*^-1^for the specific peptide and 2.5 *s*^-1^in absence of specific peptide. As expected, the non randomly distributed residues indicate that the fit does not correspond to the physical reality. The lumped off rate fit however allows to discriminate between two populations and evaluate their avidity.

## 4. Discussion

We demonstrated the development of a novel microfluidic chip capable of performing in flow interaction assays based on two distinct phenomena for the independent manipulation of two micro-sized objects. The orthogonal manipulation of the two objects allows a spatial and time control over the contact, which unlocks for the first time different adhesion assays on chip. Two different assays were performed as proof of concept to demonstrate the possibilities achievable with this method. First a biophysical experiment was performed in which fibroblasts were put in contact with fibronectin coated beads during different contact times. The adhesion mediated by the binding of integrin receptors on the surface of the fibroblasts to fibronectin was assessed and the binding kinetics of the receptor ligand pair were extracted by fitting the theoretical curve to the data. Second, human T cell clones were put in contact to cancer cells pulsed with a peptide to which the TCRs are specific. The pairs lifetime was measured after a contact time of 22 seconds, indicating a longer adhesion of T cells to cancer cells pulsed with the specific peptide than to the cancer cells not pulsed with the specific peptide. The data was approximated by a single bond dissociation to extract a pair lumped off rate describing the avidity of the interaction. This second assay opens the door to application in cancer im-munotherapy for specific T cell selection via avidity evaluation (13,28). Combination of lifetime measurement together with another criteria indicating a specific activation of the T cells, for example intracellular calcium increase (10,13) or a change of the cell’s electrical impedance (29), could allow a precise assay for specific T cell recognition. Future development will comprise the addition of a dielectrophoresis actuated cell sorter (DACS) downstream of the contact chamber to sort and retrieve cells of interest (30). However, the largest improvement will come from computer vision automation of the actuation for the reasons described below.

Reverse binding constants are known to be dependent on the force applied to disrupt the bonds. First order forward and backward kinetics were described by a single energy barrier in the potential landscape along the distance between the receptor and the ligand. The effect of a force *F* applied on the complex can be understood as a lowering in the energy barrier and an increase in the off rate, as described by the Bell model (31):

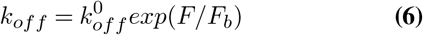

with 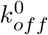 the reverse binding constant under zero force and *F_b_* the force necessary to lower the energy barrier by one unit of thermal energy *k_b_T* with *k_b_* the Boltzmann constant and *T* the absolute temperature. This was later described as a slip bond, which was verified for numerous receptor-ligand complexes (32). Integrins however were shown to display a minimum in reverse binding constant as a function of disruptive force, also known as catch bond (33). In the present case, the disruptive force is generated by the flow drag force exerted on the DMPs after the dielectrophoretic force of the contact line is turned off. Repeatable flow rates are thus of prime importance for comparable results. We ensure this by designing a serpentine that acts as large hydraulic flow resistance to minimize flow variation with pressure variation or clogs. However, feedback activated controlled particle image velocimetry (PIV) (34) would allow a dynamic and more precise control of the flow and thus of forces acting on the DMPs.

Variation of the compressive force pushing the two objects together changes the area of contact between them and has an impact on the parameter *A_c_m_min_m_max_k_on_*. The compressive force is defined by the difference between drag and DEP force and this latter is dependent on the volume of the DMPs, but also on the position of the HTPs surface. Large variation on both HTPs and DMPs size thus have an impact on both the compressive force and the disruptive force experienced by the DMPs and alter the extracted binding kinetics, which should be taken into account when working with samples of large size distribution. Using computer vision control, the voltage used to compress the DMPs in contact to the HTPs could thus be dynamically adapted to both sizes to reach constant a compressive force. Precise control on the time of contact is not possible when defining the time of contact as *t_V_sync_=0_ –t_V_contact_=0_*, indeed particles have a distribution in velocity due to differences in size and position along the channel height and do not stabilize in contact simultaneously. Control on the time of contact for less than 6 seconds was thus made a single trap at a time, but the minimum contact time was limited to 1 second due to human limitation in reactivity. This aspect would also be solved using automated control of the interaction, together with identification and sorting of events of interest.

## 5. Conclusion

In summary, we proposed and developed a new microfluidic chip combining two types of actuation for the controlled contact and separation of two micro-sized objects, in particular single-cells, and their independent controlled motions. Hydrodynamic traps were designed to first trap and arrange single-cells in the chip, and the dielectrophoresis phenomenon is used on the second type of particles, either beads or cells, to bring them towards the first cells and create a forced monitored interaction. This developed tool is specially designed to guarantee manipulation methods that preserve cells integrity and receptors functions and paves the way to a second generation of larger throughput devices as it can easily be combined with automation. We performed two different assays to demonstrate the capability of the device to generate repeatable cell-bead and cell-cell interactions, first on the fibronectin-integrin bond to fit the experimental data to the well-known binding kinetic model and validate the device. Finally, we tackled the challenge of TCR-pMHC bond and succeeded in discriminating between specific and nonspecific interactions which shows the potential of the device in cell-based cancer immunotherapy development once combined with automation for faster and less expensive T cell screening and sorting. This novel method thus opens new perspectives for applications in biophysical experiments and adoptive cell therapy developments.

## ACKNOWLEDGEMENTS

This work was supported by the EIPHI Graduate School under Contract ANR-17-EURE-0002, by the French Agence Nationale de la Recherche and the Swiss National Science Foundation through the CoDiCell project (contract “ANR-17-CE33-0009” and “FNS 00021E_175,592/1”, respectively), and by the French ROBOTEX network and its Micro and Nanorobotics center under Grant ANR-10-EQPX-44-01. The authors thank Olivier Thoumine for the fruitful discussions, Laura Collodel for her help with the experiments, François Marionnet for the valuable technical advice, and the french RENATECH network and its FEMTO-ST technological facility as well as the EPFL’s Center of MicroNanoTechnology (CMi) for the support with the microfabrication.

## Bibliography

1. Priyabrata Pattnaik. Surface plasmon resonance: Applications in understanding receptorligand interaction. Applied Biochemistry and Biotechnology, 126(2):079–092, 2005. doi: 10.1385/abab:126:2:079.

2. Xinxin Hang, Shiqi He, Zaizai Dong, Grayson Minnick, Jordan Rosenbohm, Zhou Chen, Ruiguo Yang, and Lingqian Chang. Nanosensors for single cell mechanical interrogation. Biosensors and Bioelectronics, 179:113086, May 2021. ISSN 09565663. doi: 10.1016/j.bios.2021.113086.

3. Jonne Helenius, Carl-Philipp Heisenberg, Hermann E. Gaub, and Daniel J. Muller. Singlecell force spectroscopy. Journal of Cell Science, 121(11):1785–1791, 2008. doi: 10.1242/jcs.030999.

4. Eva Potthoff, Orane Guillaume-Gentil, Dario Ossola, Jerome Polesel-Maris, Salome LeibundGut-Landmann, Tomaso Zambelli, and Julia A. Vorholt. Rapid and serial quantification of adhesion forces of yeast and mammalian cells. PLOS One, 7(12):e52712, 2012. doi: 10.1371/journal.pone.0052712.

5. Scott E. Chesla, Periasamy Selvaraj, and Cheng Zhu. Measuring two-dimensional receptorligand binding kinetics by micropipette. Biophysical Journal, 75(3):1553–1572, 1998. doi: 10.1016/s0006-3495(98)74074-3.

6. Wei Chen, Evan A. Evans, Rodger P McEver, and Cheng Zhu. Monitoring receptor-ligand interactions between surfaces by thermal fluctuations. Biophysical Journal, 94(2):694–701, 2008. doi: 10.1529/biophysj.107.117895.

7. Olivier Thoumine, Pierre Kocian, Arlette Kottelat, and Jean-Jacques Meister. Short-term binding of fibroblasts to fibronectin: optical tweezers experiments and probabilistic analysis. European Biophysics Journal, 29(6):398–408, 2000. doi: 10.1007/s002490000087.

8. Siqing Dai, Jingyu Mi, Jiazhen Dou, Hua Lu, Chen Dong, Li Ren, Rong Zhao, Wenpu Shi, Nu Zhang, Yidan Zhou, Jiwei Zhang, Jianglei Di, and Jianlin Zhao. Optical tweezers integrated surface plasmon resonance holographic microscopy for characterizing cell-substrate interactions under noninvasive optical force stimuli. Biosensors and Bioelectronics, 206: 114131, June 2022. ISSN 09565663. doi: 10.1016/j.bios.2022.114131.

9. Michael Kirschbaum, Magnus Sebastian Jaeger, Tim Schenkel, Tanja Breinig, Andreas Meyerhans, and Claus Duschl. T cell activation on a single-cell level in dielectrophoresis-based microfluidic devices. Journal of Chromatography A, 1202(1):83–89, 2008. doi: 10.1016/j.chroma.2008.06.036.

10. Burak Dura, Mariah M. Servos, Rachel M. Barry, Hidde L. Ploegh, Stephanie K. Dougan, and Joel Voldman. Longitudinal multiparameter assay of lymphocyte interactions from onset by microfluidic cell pairing and culture. Proceedings of the National Academy of Sciences, 113(26),2016. doi: 10.1073/pnas.1515364113.

11. Yufu Zhou, Ning Shao, Ricardo Bessa de Castro, Pengchao Zhang, Yuan Ma, Xin Liu, Feizhou Huang, Rong-Fu Wang, and Lidong Qin. Evaluation of single-cell cytokine secretion and cell-cell interactions with a hierarchical loading microwell chip. Cell Reports, 31(4): 107574, 2020. doi: 10.1016/j.celrep.2020.107574.

12. Patricia Moura Rosa, Nimi Gopalakrishnan, Hany Ibrahim, Markus Haug, and Oyvind Ha-laas. The intercell dynamics of T cells and dendritic cells in a lymph node-on-a-chip flow device. Lab on a Chip, 16(19):3728–3740, 2016. doi: 10.1039/c6lc00702c.

13. Julian F. Ashby, Julien Schmidt, Neelima KC, Armand Kurum, Caroline Koch, Alexandre Harari, Li Tang, and Sam H. Au. Microfluidic T cell selection by cellular avidity. Advanced Healthcare Materials, 11(16):2200169, 2022. doi: 10.1002/adhm.202200169.

14. Margaux Duchamp, Thamani Dahoun, Clarisse Vaillier, Marion Arnaud, Sara Bobisse, George Coukos, Alexandre Harari, and Philippe Renaud. Microfluidic device performing on flow study of serial cell–cell interactions of two cell populations. RSC Advances, 9(70): 41066–41073, 2019. doi: 10.1039/c9ra09504g.

15. Max A. Stockslager, Josephine Shaw Bagnall, Vivian C. Hecht, Kevin Hu, Edgar Aranda-Michel, Kristofor Payer, Robert J. Kimmerling, and Scott R. Manalis. Microfluidic platform for characterizing TCR-pMHC interactions. Biomicrofluidics, 11(6):064103, 2017. doi: 10.1063/1.5002116.

16. Hiroki Ide, Wilfred Villariza Espulgar, Masato Saito, Taiki Aoshi, Shohei Koyama, Hy-ota Takamatsu, and Eiichi Tamiya. Profiling T cell interaction and activation through microfluidics-assisted serial encounter with APCs. Sensors and Actuators B: Chemical, 330:129306, 2020. doi: 10.1016/j.snb.2020.129306.

17. Steven A. Rosenberg and Nicholas P. Restifo. Adoptive cell transfer as personalized im-munotherapy for human cancer. Science, 348(6230):62–68, 2015. doi: 10.1126/science.aaa4967.

18. Clémentine Lipp, Kevin Uning, Jonathan Cottet, Daniel Migliozzi, Arnaud Bertsch, and Philippe Renaud. Planar hydrodynamic traps and buried channels for bead and cell trapping and releasing. Lab on a Chip, 21(19):3686–3694,2021. doi: 10.1039/d1lc00463h.

19. Clémentine Lipp, Laure Koebel, Arnaud Bertsch, Michaёl Gauthier, Aude Bolopion, and Philippe Renaud. Dielectrophoretic traps for efficient bead and cell trapping and formation of aggregates of controlled size and composition. Frontiers in Bioengineering and Biotechnology, 10, 2022. doi: 10.3389/fbioe.2022.910578.

20. Ying Zhou, Srinjan Basu, Ernest Laue, and Ashwin A. Seshia. Single cell studies of mouse embryonic stem cell (mESC) differentiation by electrical impedance measurements in a mi-crofluidic device. Biosensors and Bioelectronics, 81:249–258, July 2016. ISSN 09565663. doi: 10.1016/j.bios.2016.02.069.

21. Srinivasu Valagerahally Puttaswamy, Nikhil Bhalla, Colin Kelsey, Gennady Lubarsky, Chengkuo Lee, and James McLaughlin. Independent and grouped 3D cell rotation in a microfluidic device for bioimaging applications. Biosensors and Bioelectronics, 170:112661, December 2020. ISSN 09565663. doi: 10.1016/j.bios.2020.112661.

22. Nicolas Demierre, Thomas Braschler, Pontus Linderholm, Urban Seger, Harald van Lintel, and Philippe Renaud. Characterization and optimization of liquid electrodes for lateral dielectrophoresis. Lab on a Chip, 7(3):355–365, 2007. doi: 10.1039/b612866a.

23. A Ramos, H Morgan, N G Green, and A Castellanos. Ac electrokinetics: a review of forces in microelectrode structures. Journal of Physics D: Applied Physics, 31(18):2338–2353, 1998. ISSN 0022-3727, 1361–6463, doi: 10.1088/0022-3727/31/18/021.

24. Jonathan Cottet, Alexandre Kehren, Soufian Lasli, Harald Lintel, Francois Buret, Marie Frenea-Robin, and Philippe Renaud. Dielectrophoresis-assisted creation of cell aggregates under flow conditions using planar electrodes. ELECTROPHORESIS, 40(10):1498–1509, 2019. doi: 10.1002/elps.201800435.

25. Jeong-Ryul Hwang, Yeongseon Byeon, Donghwan Kim, and Sung-Gyoo Park. Recent insights of T cell receptor-mediated signaling pathways for T cell activation and de-velopment. Experimental & Molecular Medicine, 52(5):750–761, 2020. doi: 10.1038/s12276-020-0435-8.

26. Johannes B. Huppa and Mark M. Davis. T-cell-antigen recognition and the immunological synapse. Nature Reviews Immunology, 3(12):973–983, 2003. doi: 10.1038/nri1245.

27. Diana Campillo-Davo, Donovan Flumens, and Eva Lion. The quest for the best: How TCR affinity, avidity, and functional avidity affect TCR-engineered T-cell antitumor responses. Cells, 9(7):1720, 2020. doi: 10.3390/cells9071720.

28. Taylor Yerbury. z-Movi cell avidity analyzer - LUMICKS, 2022.

29. Enrica Rollo, Enrico Tenaglia, Raphaël Genolet, Elena Bianchi, Alexandre Harari, George Coukos, and Carlotta Guiducci. Label-free identification of activated T lymphocytes through tridimensional microsensors on chip. Biosensors and Bioelectronics, 94:193–199, August 2017. ISSN 09565663. doi: 10.1016/j.bios.2017.02.047.

30. Dongkyu Lee, Bohyun Hwang, and Byungkyu Kim. The potential of a dielectrophoresis activated cell sorter (DACS) as a next generation cell sorter. Micro and Nano Systems Letters, 4(1),2016. doi: 10.1186/s40486-016-0028-4.

31. George I. Bell. Models for the specific adhesion of cells to cells. Science, 200(4342):618–627, 1978. doi: 10.1126/science.347575.

32. Baoyu Liu, Wei Chen, and Cheng Zhu. Molecular force spectroscopy on cells. Annual Review of Physical Chemistry, 66(1):427–451, 2015. doi: 10.1146/annurev-physchem-040214-121742.

33. Fang Kong, Andrés J. Garcia, A. Paul Mould, Martin J. Humphries, and Cheng Zhu. Demonstration of catch bonds between an integrin and its ligand. Journal of Cell Biology, 185(7): 1275–1284, 2009. doi: 10.1083/jcb.200810002.

34. Stefan Kobel, Ana Valero, Jonas Latt, Philippe Renaud, and Matthias Lutolf. Optimization of microfluidic single cell trapping for long-term on-chip culture. Lab on a Chip, 10(7):857, 2010. doi: 10.1039/b918055a.

