## Supplementary Figures for "Microfluidic device combining hydrodynamic and dielectrophoretic trapping for the controlled contact between single micro-sized objects and application to adhesion assays"

### Supplementary material

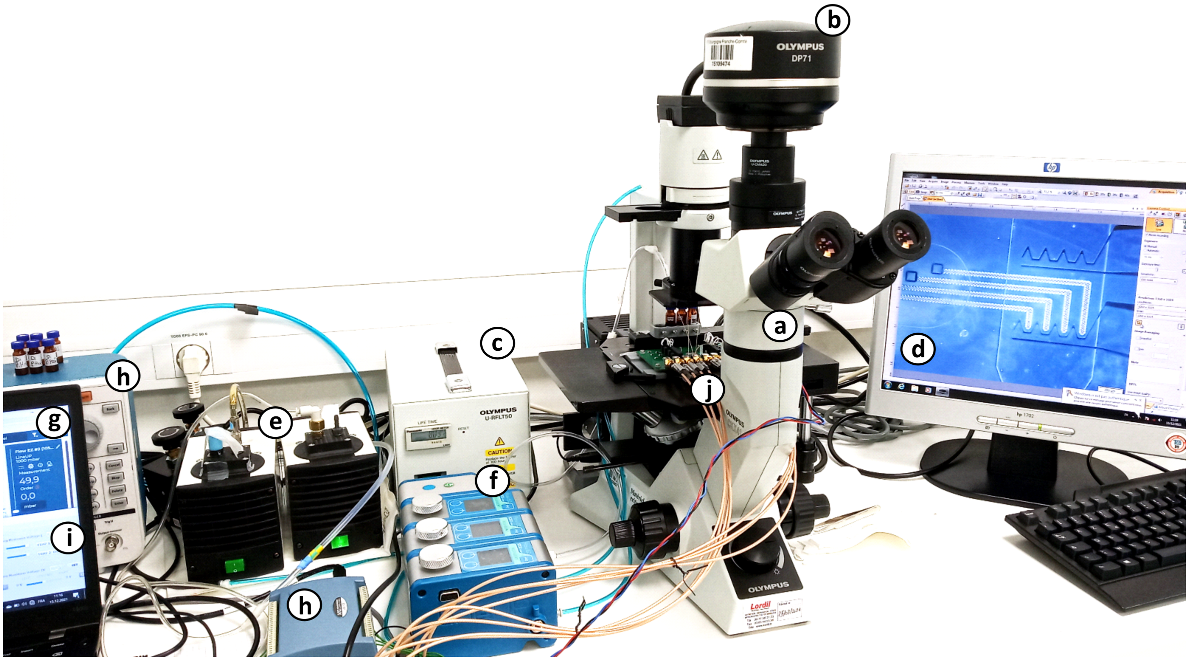


**Figure S 1:** Picture of the experimental set-up, composed of the imaging system (**a-d**) , the fluidic system (**e-g**) and the dielectrophoretic actuation system (**h-j**). The imaging system comprises a fluorescence microscope (**a**), a camera (**b**), a fluorescence light source (**c**) and the software to monitor the cells interacting in the device (**d**). The fluidic system consists in air pumps (**e**) generating negative and positive pressure outlets for the pressure controller (**f**) to deliver the right controlled pressures to the fluid tanks, asked by the pressure controller software (**g**). Finally, to generate the dielectrophoresis in the chip, electronic signals are produced by a custom PCB multiplying an AC signal from a signal generator with DC signals (**h**) whose different amplitudes are set on a custom C++ controlling software (**i**). The software enables the control of the trajectory of each single cell, and the duration of the interactions. All of the generated signals are then sent to another custom PCB on which the chip is mounted (**j**).

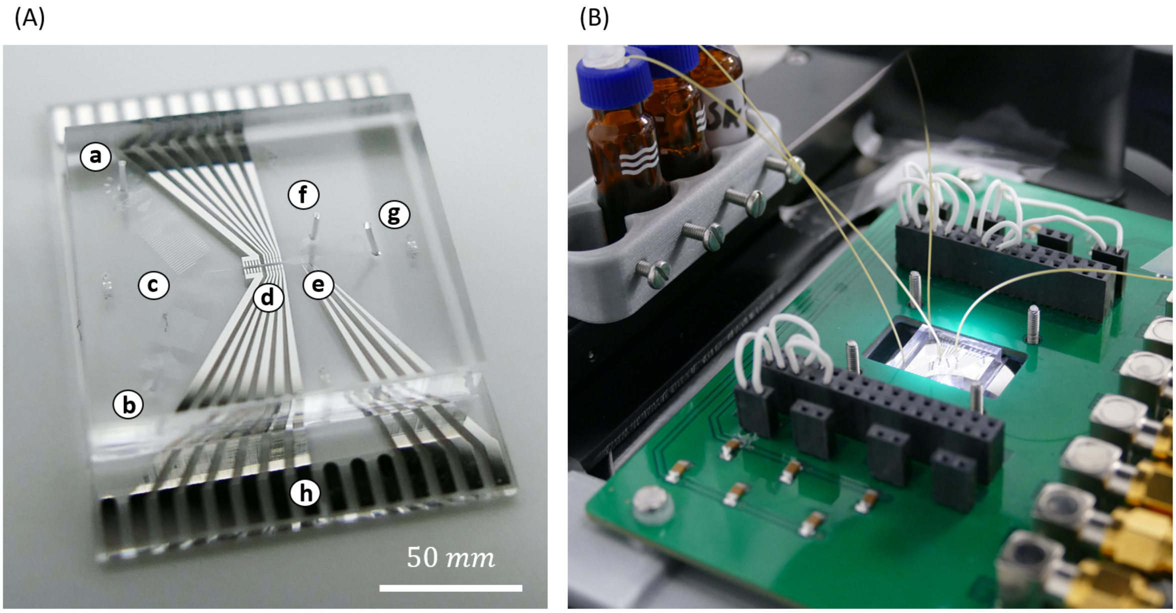


**Figure S 2:** **(A)** Picture of a chip illustrated in Figure 1 a with two inlets (**a**-**b**) and corresponding serpentine channels (**c**) that merge before entering the deviation zone (**d**), followed by the interaction chamber (**e**), the hydrodynamic trapping in/outlet on its side (**f**) and the main outlet (**g**). The electrodes are also visible with respective connecting pads on the side (**h**). **(B)** Picture of the chip mounted on the PCB for electrical connection. The pressures in the vials are controlled via pressure pumps and a tubing plunged in the liquid conducts the fluid to the chip.

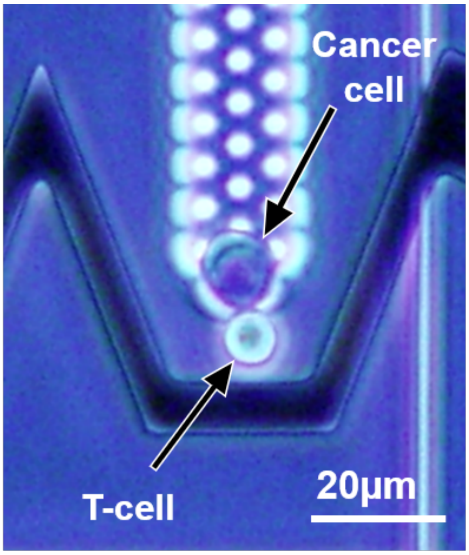


**Figure S 3:** Optical microscope picture of a T cell forced in contact to a cancer cell. The side of the PDMS channel is visible on the right and shows the good alignment between top PDMS channel and the electrodes. The buried channel is especially bright because of its uneven topography that reflects light.

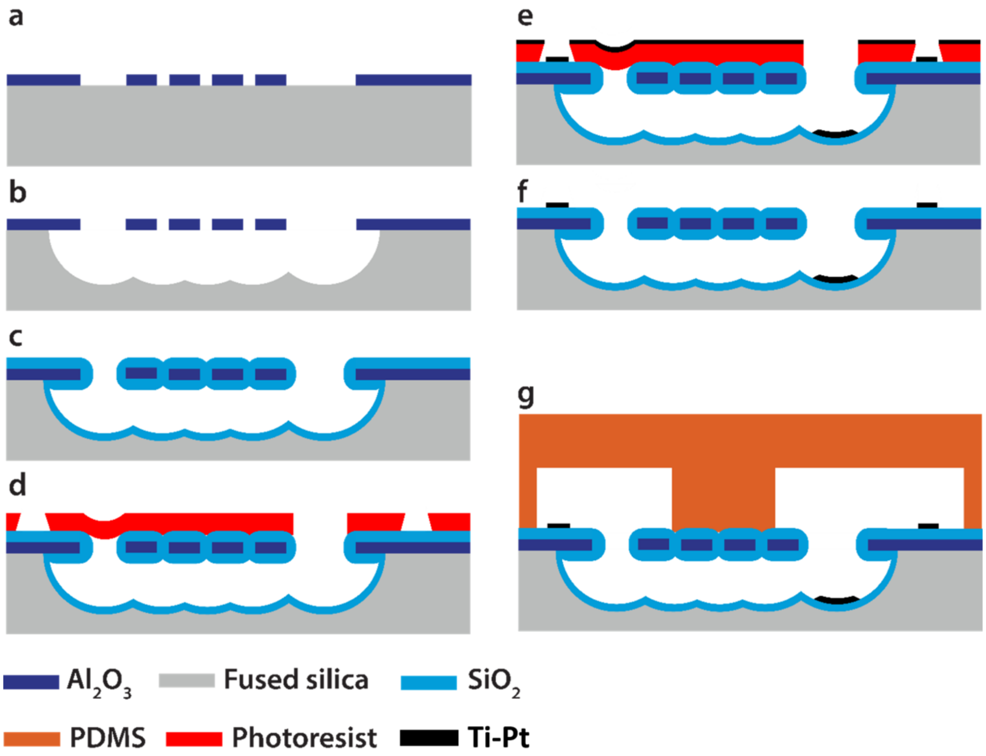


**Figure S 4:** Process for the fabrication of buried channels with electrodes: **a** deposition and patterning of a 300 nm thick aluminum oxide thin film. **b** Selective underetching of the fused silica substrate in vapor HF. **c** Deposition of 2.5 μm of low temperature oxide (silicon dioxide) to close the access holes but leaves open the traps and outlets. **d** Photolithography patterning of a lift-off resist with undercut. The photoresist covers the small openings like the traps thanks to surface tension of the resist but does not fill the large ones. **e** Evaporation of 20 nm - 200 nm Ti-Pt thin films. **f** Lift-off process, the metal remains in the large opening where it is not problematic but is lifted from the small ones where transparency is needed. **g** Alignment and permanent bonding of the top PDMS channels.

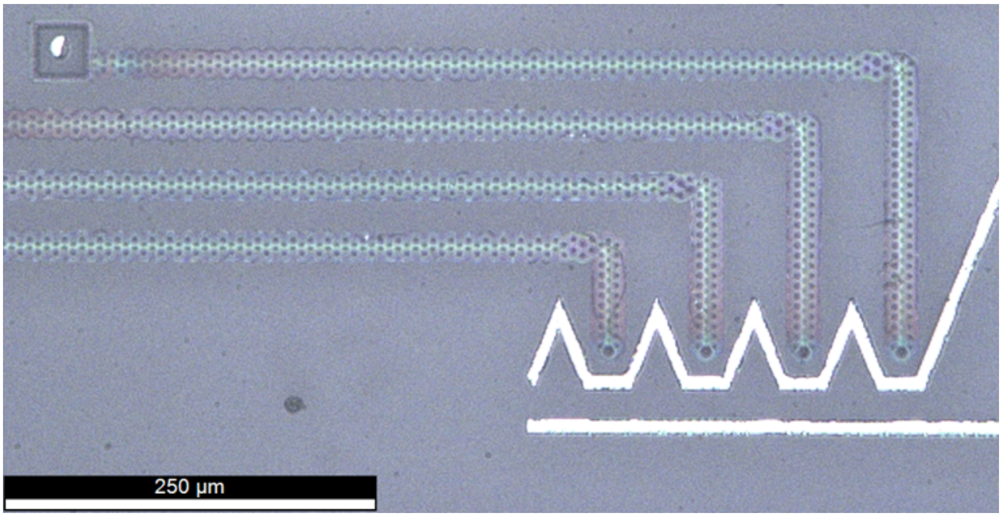


**Figure S 5:** Optical microscope picture of the buried channels and electrodes after the lift-off process: metal can be observed in the outlets of the buried channels (top left) but the hydrodynamic traps are free from metal and have good transparency.

| | | 0 hours | 5 hours | 24 hours | 48 hours |
|---|---|---|---|---|---|
| **Colo205** | Complete medium (all times) | 50 000 cells | 52 000 cells | 132 000 cells | 216 000 cells |
| | 10% PBS medium (5 h) then complete medium (24 and 48 h) | 50 000 cells | 10 000 cells | 0 cells | 0 cells |
| **CD8 T cells** | Complete medium (all times) | 100 000 cells | 96 000 cells | 80 000 cells | 52 000 cells |
| | 10% PBS medium (5 h) then complete medium (24 and 48 h) | 100 000 cells | 56 000 cells | 8 000 cells | 8 000 cells |

**Figure S 6:** Viability of Colo205 and CD8 T cells at different stages after 5h immersion in 10% PBS diluted in DI water complemented in dextrose for osmolarity compared to complete medium (DMEM and RPMI respectively). After the 5 h incubation in the 10% PBS medium, cells were washed and cultured in standard medium. The 5h in the 10% PBS medium is incompatible with the viability of both types of cells as it drops drastically in both cases even when the cells are put back into their original complete medium after the 5h. A further study on medium dilutions is thus needed to obtain the maximum dilution possible keeping a high viability of cells (see Supplementary Figure S7).

| | 0 hours | 5 hours | 24 hours | 48 hours |
|---|---|---|---|---|
| **Complete medium (100%)** | 500 000 cells (100% cell viability) | 400 000 cells (100% cell viability) | 500 000 cells (96% cell viability) | 620 000 cells (98% cell viability) |
| **Complete medium (80%)** | 500 000 cells (100% cell viability) | 400 000 cells (100% cell viability) | 340 000 cells (97% cell viability) | 350 000 cells (96% cell viability) |
| **Complete medium (60%)** | 500 000 cells (100% cell viability) | 350 000 cells (100% cell viability) | 300 000 cells (91% cell viability) | 360 000 cells (92% cell viability) |
| **Complete medium (40%)** | 500 000 cells (100% cell viability) | 325 000 cells (97% cell viability) | 330 000 cells (89% cell viability) | 320 000 cells (87% cell viability) |
| **Complete medium (20%)** | 500 000 cells (100% cell viability) | 400 000 cells (96% cell viability) | 230 000 cells (89% cell viability) | 210 000 cells (84% cell viability) |
| **Complete medium (10%)** | 500 000 cells (100% cell viability) | 250 000 cells (93% cell viability) | 170 000 cells (85% cell viability) | 160 000 cells (89% cell viability) |

**Figure S 7:** T cell viability at different stages after 5 h immersion in complete medium diluted in different percentage of DI water with corresponding amount of dextrose to compensate for osmolarity. After the 5 h incubation in custom medium, cells were washed and cultured in standard medium.

**Video S 1:** Video of the controlled interaction between CD8 T-cell clones and Colo205 cancer cells pulsed with a peptide to which the T-cell receptors (TCR) are specific. The fluid constantly flowing from top to bottom is dragging the T-cells that are first held in the DEP synchronization line before releasing them for a synchronized departure. The cells are dragged to the contact line and arrive in contact with the cancer cells previously placed in the hydrodynamic traps. The DEP contact line pushes each T-cell against each cancer cell in a forced interaction during a controlled contact duration, until the DEP traps are turned off. The T-cells are dragged away by the fluid flow and monitoring the time between the disabling of the traps and the detachment of the T-cell allows to measure the cell avidity of each interaction, and gives the possibility to discriminate between specific and nonspecific interactions.
